# Hybridization Mediated Range Expansion and Climate Change Resilience in Two Keystone Tree Species of Boreal Forests

**DOI:** 10.1101/2023.01.31.526517

**Authors:** Piyal Karunarathne, Qiujie Zhou, Martin Lascoux, Pascal Milesi

**Affiliations:** Plant Ecology and Evolution, Department of Ecology and Genetics, Uppsala University, Norbyvägen 18D, 75236 Uppsala, Sweden; Science for Life Laboratory (SciLifeLab), Uppsala, Sweden; Institute of Population Genetics, Heinrich-Heine University, Düsseldorf, Universitäts Straße 1, 40225 Düsseldorf, Germany

**Keywords:** Eco-evolution, Climate change, Landscape genomics, hybridization, forest trees

## Abstract

Current global climate change is expected to affect biodiversity negatively at all scales leading to mass biodiversity loss. Many studies have shown that the distribution of allele frequencies across a species’ range is often influenced by specific genetic loci associated with local environmental variables. This association reflects local adaptation and allele changes at those loci could thereby contribute to the evolutionary response to climate change. However, predicting how species will adapt to climate change from this type of data alone remains challenging. In the present study, we combined exome capture sequences and environmental niche reconstruction, to test multiple methods for assessing local adaptation and climate resilience in two widely distributed conifers, Norway spruce and Siberian spruce. Both species are keystone species of the boreal forest and share a vast hybrid zone. We show that local adaptation in conifers can be detected through allele frequency variation, population-level ecological preferences, and historical niche movement. Moreover, we integrated genetic and ecological information into genetic offset predictive models to show that hybridization plays a central role in expanding the niche breadth of the two conifer species and may help both species to cope better with future changing climates. This joint genetic and ecological analysis also identified genetically isolated populations that are at risk under current climate change.

## Background

Human activities have led to the destruction of the habitat of an increasing number of species and climate change has only exacerbated this situation (Bellard et al., 2012; Grimm et al., 2013; Walther et al., 2002). Given the rapid pace of global warming, ecological communities face an urgent need for rapid adaptation in their structure and composition to cope with these ongoing challenges. Interestingly, recent studies have revealed that communities tend to respond more strongly to regional than global changes (Altieri & Gedan, 2015; Araújo & Rahbek, 2006; Atkins & Travis, 2010; Thuiller et al., 2005; Williams et al., 2021; Wuethrich, 2000). Climate change impacts vary across ecosystems, leading to diverse ecological dynamics and unpredictable changes within local environments (Carbeck et al., 2022; Walther et al., 2002). In order to mitigate the risk of demographic collapse and potential extinction, organisms must either track their climatic niche, be phenotypically plastic, or adapt to the changing environment from existing genetic variation or through de novo mutations. Adaptations of populations to local conditions result from a subtle balance between the intensity of natural selection and migration between the populations. While the former pushes the populations towards their local optima the latter tends to homogenize the gene pools between the populations (for a review see Bonnet et al., 2022; Lenormand, 2002). Still, some level of gene flow, although not an absolute requirement, contributes to promoting local and global genetic diversity and preserving the adaptive potential of populations (e.g., Frankham, 2005; Markert et al., 2010; Pauls et al., 2013). Global climate change poses significant challenges as it can affect simultaneously many populations, placing species and their populations at greater vulnerability in limiting their ability to cope with environmental changes and potentially leading to extinction (Leroy et al., 2020; Pauls et al., 2013; Rellstab et al., 2016). Plants are probably even more affected by global change than animals due to their inherent disadvantage in tracing suitable habitats, especially species with long generation times (Dauphin et al., 2021; Leroy et al., 2020). Therefore, the survival of a long-lived plant species depends critically upon the potential of the species to expand into new regions through past adaptations to such environments as well as the potential for populations at the edge of its distribution range to face new environmental conditions (Pfennig et al., 2016; Taylor et al., 2015). Therefore, to comprehensively understand the interplay between environmental shifts and their genetic consequences, it is imperative to employ eco-evolutionary models (Bush et al., 2016; Thuiller et al., 2013). For instance, Cotto et al., (2017) pioneered an innovative approach by integrating ecological niche modeling with individual-based genetics modeling to explore how populations in the Alpine range respond to evolving environmental conditions. Similarly, genetic offset methodologies are designed to predict the disparity between a population’s current status and the necessary adaptations to be able to survive under future climatic scenarios (e.g., Rellstab et al., 2021).

However, these approaches encounter certain limitations when applied on a continental scale, particularly with regard to accounting for intricate demographic histories and ongoing gene flow dynamics. In our present study, we have taken a multifaceted approach by seamlessly merging ecological niche modeling, landscape genomics, and genotype-environment association analyses. Our aim is to unravel the pivotal role of recurrent hybridization in facilitating the adaptation of two cornerstone species within the boreal forest ecosystem to the challenges posed by global environmental changes.

In Western Europe, recurrent cycles of population contraction and expansion following glacial cycles shaped both genetic diversity (Milesi et al., 2023; Petit et al., 2003) and local adaptation patterns along environmental gradients that also correspond to post-glacial recolonization routes (L. Li et al., 2022; Savolainen et al., 2013; Thörn et al., 2021). Genetic admixture following range expansion from refugia can serve as a major source of genetic diversity in forest trees (J. Chen et al., 2019; Lexer et al., 2010; Müller et al., 2019; Tsuda et al., 2017; Zhou et al., 2023). It can promote increased climate resilience in wild populations (e.g., Inoue & Berg, 2017; Meester et al., 2018) and aid wider phenotypic plasticity against rapid climate changes (see Catullo et al., 2019)). Admixture caused by recurrent colonization by superior ecotypes can promote better resistance to environmental stress (e.g., (Hamilton & Miller, 2016; Ortego et al., 2015) notably through an effective population size increase, persistence at the range edge and niche expansion (e.g., Gillies et al., 2016; Teukeng et al., 2022; Zalapa et al., 2010). Hybridization and admixture are thus effective mechanisms to increase overall genetic diversity and can further promote the transfer of adaptive alleles from one genetic background to another, enhancing local adaption and allowing populations to respond quickly to selective pressures (Rendón-Anaya et al., 2021; Rieseberg & Willis, 2007; Stelkens et al., 2014; Taylor et al., 2015). Here we hypothesize that the recurrent hybridization and admixture events between two closely related coniferous species through cycles of historic range shifts due to ice cycles may help both species to better cope with future climatic changes.

Norway spruce (*Picea abies* (L.) H. Karst) and Siberian spruce (*Picea obovata* Ledeb.) are two closely related conifer species that diverged several million years ago (J. Chen et al., 2019; Tsuda et al., 2016; Zhou et al., 2023). Today, the two species have clear western (Western Europe) and eastern (Siberian Russia) distributions, albeit with a known hybrid zone located between west of the Ural Mountains and expanding up to Northern Fennoscandia (Tollefsrud et al., 2015; Tsuda et al., 2016; Zhou et al., 2023: See map Fig. 1). Currently, the distribution of *P. abies* is subdivided into seven main genetic clusters (J. Chen et al., 2019; L. Li et al., 2022). They correspond to three ancestral clusters (Alpine, Carpathian, Fennoscandian) as well as to admixture between these three main clusters (J. Chen et al., 2019; L. Li et al., 2022). Siberian spruce, on the other hand, expanded westwards from Northeast Asia to Western Russia. Following glacial cycles, the two species came into contact on several occasions, leading to introgression (e.g., J. Chen et al., 2019; Tollefsrud et al., 2015; Tsuda et al., 2016; Zhou et al., 2023). Recently, Nota et al., (2022), using ancient DNA, and Li et al., (2022), using contemporary nuclear markers, confirmed conclusions drawn from pollen records (e.g. Giesecke, 2005) that recolonization of the Scandinavian peninsula by Norway spruce after the last glacial maximum (LGM) occurred through two different entry points, a southern one, from Eastern location across the Baltic Sea and a northern one through northern Finland with a contribution from *P. obovata*. Li et al. (2022) demonstrated that the contact zone between the two main genetic clusters aligns with the boundary between Scandinavia’s primary climatic regions, revealing the influence of natural selection in maintaining this zone and shaping spruce genetic diversity. While pollen records are valuable for inferring species distribution and population movement during the Holocene (e.g., Trondman et al. 2014), they offer limited insights, particularly at the species level, for events predating that period, as observed in the case of Norway and Siberian spruce (J. Chen et al., 2019; Tsuda et al., 2016; Zhou et al., 2023).

**Figure 1:**
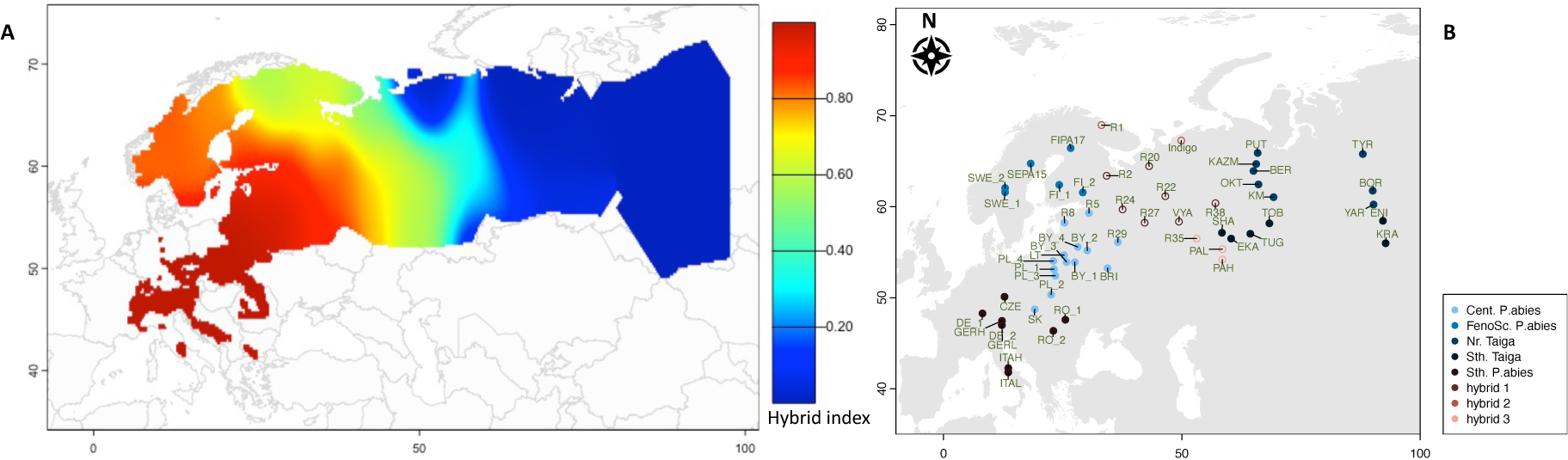
Climatic clusters and hybrid index (*hi*) between *P. abies* and *P. obovata;* **A.** Spatial projection of the hybrid index computed from ancestry coefficient using K=2 [For the analyzed alleles, *hi =* 1 = *P. abies* = red, *hi =* 0 = *P.obovata* = blue]. **B.** Genetically informed climatic clusters of the studied populations; clusters were identified with k-means clustering of environmental and adaptive index variables (see methods) [Cent: central, FenoSc: Fennoscandian, Nr: northern, Sth: southern].

We here used a combination of genome scans, genotype-environment association studies, and ecological niche modeling (for both past and future scenarios) to demonstrate that recurrent hybridization significantly contributes to the expansion of both species’ niche breadth, potentially enabling them to better cope with changing climates. Given the close genetic affinity, the extensive hybrid zone, historic recolonization events, and the climatic gradient occupied by Norway and Siberian spruce, they provide a unique system for studying the role of hybridization in shaping climate resilience through genetic variation and range expansion. Our study first provides evidence of local adaptation and identifies key climatic variables influencing species distributions. We then use environmental niche models with these climatic variables and species occurrences to define the ecological niche preferences of both species and their historical movements. Finally, we integrate genetic and ecological information into genetic offset predictive models to assess the resilience of each population to future climatic conditions.

## Results

### Populations exhibiting a high level of admixture are spread across three climatic groups

We used the same sampling of 55 populations covering the whole range of *P. abies* and the western part of the distribution range of *P. obovata* as in Zhou et al. (2023). Using a combination of population genetics analyses, the authors showed that there are two main genetic entities: one western cluster, corresponding to Norway spruce (*P. abies*), centered on the Alps and the Carpathians, and extending towards Scandinavia and the Russian Plains, and one eastern cluster stemming from Siberia, east of the Urals Mountains and corresponding to what has generally been recognized as Siberian spruce (*P. obovata*). The populations previously recognized as of hybrid nature located west of the Ural Mountains and east of the Baltic region formed sub-clusters within the two main clusters (Fig. S1A also see Zhou et al. 2023 Fig. S3). Following this initial clustering, in order to quantify the degree of admixture between *P. abies* and *P. obovata*, we derived a *hybrid index* for each individual using ADMIXTURE (K=2, see materials and methods and Fig. 1A & S1B) and averaged it per population. The distribution of the *hybrid index* is highly consistent with the Bayesian clustering. To investigate whether specific DNA polymorphisms were associated with given climatic zones, we used redundancy analysis (RDA) incorporating both genetic and climatic data (see *Genetic diversity distribution among populations* in materials and methods). We defined five major environmental clusters (i.e., genetically informed climatic groups) among all the populations studied, with populations in the hybrid zone belonging to three different clusters (Fig. 1B). Interestingly, this divided western populations (i.e., *P. abies*) into three groups: a southern group consisting of Alpine, Carpathian and Central European populations, a northern group made of Fennoscandian populations and, a third group with central European and Baltic populations, while eastern populations (i.e., *P. obovata*) were separated into a northern and a southern group along the southern border of the west Siberian taiga (Fig. 1B).

### Spatial heterogeneity and environmental conditioning of genetic diversity

We investigated changes in total nucleotide diversity (π) across the whole distribution range of the species. Globally, π ranging from 0.00518 to 0.00695, the highest values were found in the North-Eastern range for *P. obovata* and Southern Scandinavia for *P. abies*. Overall, *P. obovata* populations exhibit higher π values than *P. abies*, likely reflecting more stable historical distribution and more continuous distribution through space (Fig. 2A). In *P. obovata*, a clear shift in π matches the location of the Ural Mountain, which likely limits the gene flow. Pockets of high diversity are found in the Southernmost range of *P. obovata*, west of the Urals, possibly reflecting refugia during the ice age. While the overall genetic diversity within *P. abies* also ranges between 0.0054 and 0.0068, the northern- and southern-most populations had the lowest nucleotide diversity. Even though southern locations in Italy and the Alpine mountains region have been refugia for Norway spruce during the ice ages, the current relative isolation of the populations because of the topography, probably explains the relatively lower genetic diversity. On the other hand, the northernmost populations of *P. abies* in Sweden harbor the lowest genetic diversity. These populations are also the youngest and the low nucleotide diversity may reflect the colonization forefront. Nevertheless, the highest π values are found in Southern Scandinavia, an area that was also recolonized after the LGM but from different populations. Finally, populations located in contact zones (e.g., in Scandinavia between the Northern and the Southern range) or located within the large hybrid zone between *P. abies* and *P. obovata* exhibit higher π values than neighboring populations, exhibiting the effect of introgression and/or admixture on genetic diversity. As expected, hybrid populations also show intermediate heterozygosity between *P. abies* and *P. obovata* (linear model, *F* = 6.55, *df* = 34, *p* = 0.015, Fig. 2B). More importantly, hybrid populations tend to have more homogeneous allele frequencies from one SNP to another than in *P. abies* (Welch *t*-test, *t* = -2.85, *df* = 11.172, *p* = 0.02) or *P. obovata* (*t* = -5.63, *df* = 7.93, *p* < 0.001, Fig. 2C): Note that standard deviation in allele frequencies vary quantitively with the hybrid index, meaning that they are both minimum for a hybrid index closed to 0.5 (Fig. S4 A&B).

**Figure 2.**
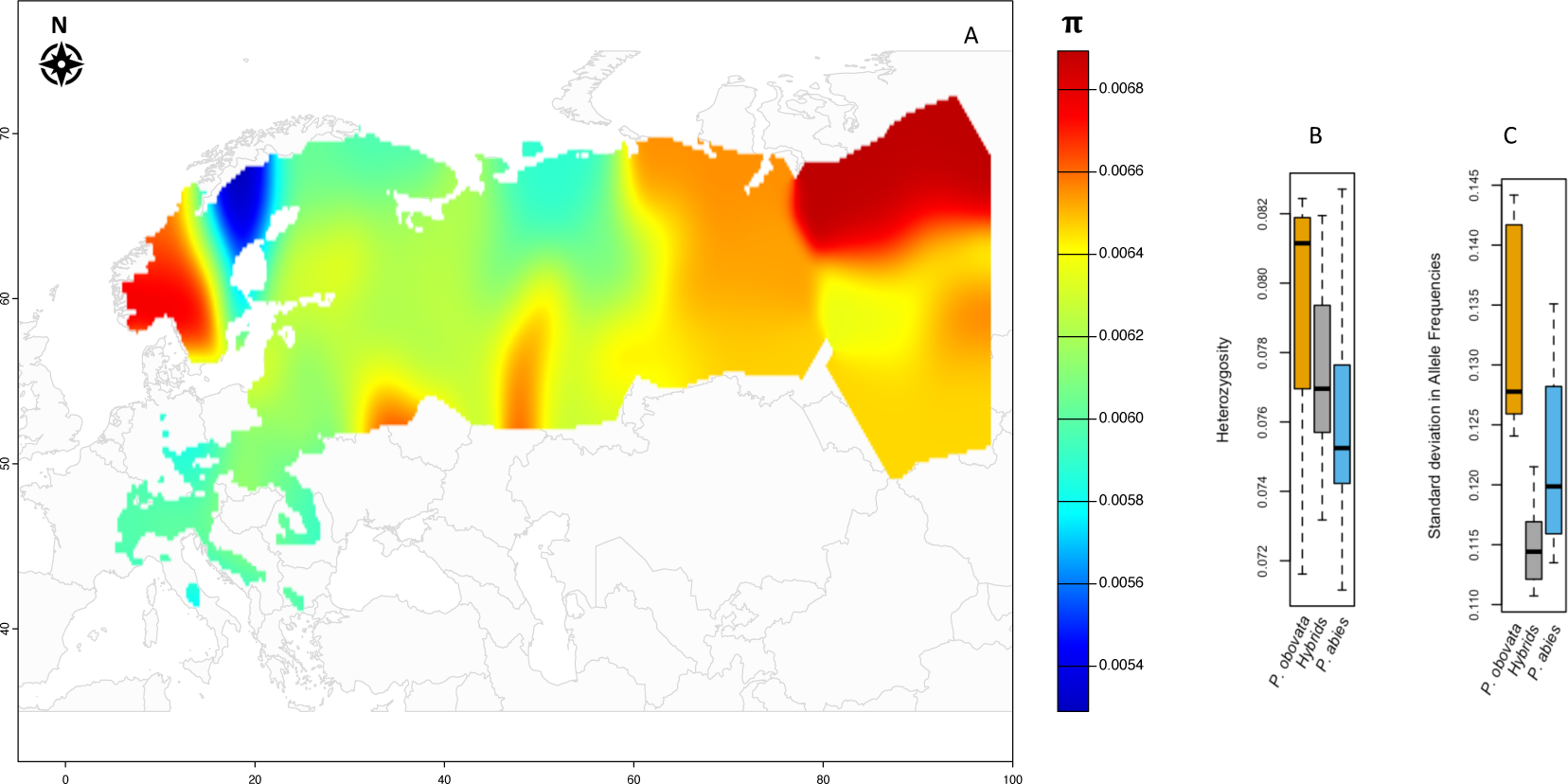
**A.** The distribution of nucleotide diversity (π) throughout the complete distribution range of the two species. **B.**, and **C.** depict population-wise heterozygosity and average and standard deviation in allele frequencies across SNPs for *P. obovata*, *P. abies,* or hybrid populations.

### Local adaptation

We further investigated whether the observed pattern of genetic differentiation could be due to local adaptations using *pcAdapt*, a genome scan method to detect outlier SNPs exhibiting a higher degree of differentiation than the genetic background while controlling for population structure. We detected 1650 SNPs, clustered into specific areas of the genome (Fig. S3 A&C) suggesting that natural selection has shaped the current genetic diversity, besides the population movements and demographic history. In order to understand the role of environmental variation in shaping genetic diversity, we conducted a redundancy analysis (RDA) with bioclimatic factors as predictors (Fig. S3B). After an ad-hoc selection to remove high levels of correlation between climatic variables, we retained three climatic variables: *Bio8*, the mean temperature of the wettest quarter, *Bio9*, the temperature of the driest quarter, and *Bio18*, the precipitation of the warmest quarter. While *Bio8* mainly varies along latitudes, *Bio9* and *Bio18* mainly reflect continentality and vary longitudinally. Together the climatic variables explained 5.4% of the total genetic variation in the climatic partial RDA (pRDA) where co-variables were introduced to control for population structure (Fig. S3B). Still, we detected 1536 outlier loci significantly associated with at least one of the bioclimatic variables; out of which 298 were recognized as top outliers, based on the overlap of a simple RDA model and the climatic model (see Capblancq & Forester, 2021 for details). These top outliers clustered together in various regions of the genome (Fig. S3A). Outlier SNPs showed substantial congruence with the enrichment observed in the *pcAdapt* output (82 % overlap, Fig. S3D) indicating that a significant part of the allele frequency variation observed among populations is driven by environmental heterogeneity. From the pRDA, we derived an *adaptive index* (see materials and methods) that measures the match between the genotypes and their environment to capture the spatial distribution of population-level adaptation to current climatic conditions (Fig. 3). The adaptive index and the hybrid index were highly correlated (Spearman’s rho = 0.90, S = 2156, *p* < 0.001 and rho = 0.40, S= 13339, *p* = 0.003, for adaptive index computed along RDA1 and 2 respectively). While the populations at both extremes of the sampling range show the highest degree of adaptation (strongly positive or negative values), populations located within the hybrid zone exhibit intermediate values indicating a lower sensitivity to environmental constraints (Fig. 3A & B).

**Figure 3.**
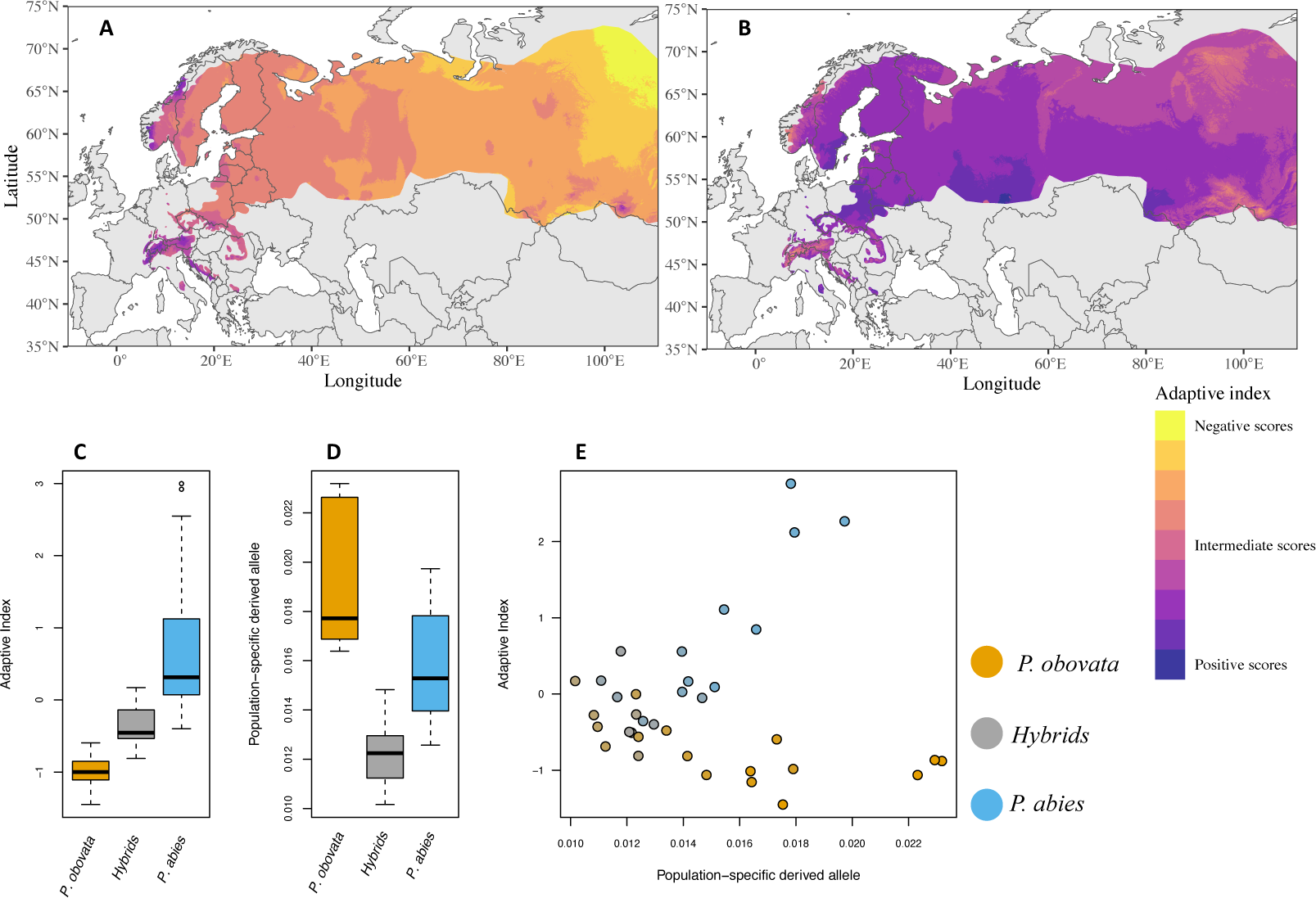
The spatial distribution of population adaptedness in Norway and Siberian spruces – the adaptive index calculated for A – RDA1 & B – RDA2. Both negative and positive adaptive index scores depict the adapteness on a continuous spectrum of opposite climatic conditions (Capblancq et al. 2021). **C.** Distribution of the adaptive index between *P. abies*, *P. obovata,* and their hybrids. **D.** Distribution of the proportion of population-specific major alleles between *P. abies*, *P. obovata,* and their hybrids **E.** Adaptive index as a function of the proportion of population-specific major alleles, color gradient represents the hybrid index for each group as in **C&D**.

We subsequently tested whether the observed pattern is primarily influenced by candidate SNPs specific to the environmental variables included in the pRDA or if it demonstrates a broader, overarching trend. It is worth noting that the phenomenon of adaptation across environmental gradients typically involves a polygenic basis, wherein alterations in allele frequencies across multiple genetic loci contribute to the observed effects. Consequently, our hypothesis anticipates that populations demonstrating heightened adaptation will exhibit distinct allele frequencies when contrasted with the wider global dataset. For each population, we calculated the ratio of major alleles specific to that population. For individual SNPs, we initially identified the minor allele as the one with the lowest frequency across the entire dataset (*f_all_* < 0.5). Subsequently, we determined, for each population individually, the percentage of sites at which the minor allele’s frequency exceeded 0.5. In other words, while the allele might be minor in the broader global context (frequency < 0.5), it assumed a major role within the focal population (i.e. *f_all_* < 0.5 but *f_popi_* > 0.5). First, the hybrid populations showed a significantly lower proportion of major alleles than that in *P. abies* or *P. obovata* populations (linear model, *r^2^* = 0.65, *df* = 33, *p* < 0.001, Fig. 3C and Fig. S4C). Second, a strong positive correlation exists between the absolute adaptive index of a population and the proportion of major alleles in that population (Spearman’s *rho* = 0.67, S= 2532, *p* < 0.001, also see Fig. 3E); the same relationship but weaker is observed when computing the adaptive index along RDA2 (Spearman’s *rho* = 0.33, S= 2502, *p* = 0.05). The reduced adaptiveness observed in hybrid populations is not solely attributed to markers associated with the environmental variables considered in the pRDA analysis. Instead, it is indicative of widespread shifts in allele frequencies throughout their genome. Additionally, *P. obovata* populations, characterized by a greater number of population-specific major alleles, tend to exhibit higher genetic diversity. Therefore, it is probable that the observed pattern is a result of selection pressures rather than genetic drift, especially in populations that may be somewhat isolated.

### Environmental preference, historical niche movement, and genetic refugia

Using the ordination of climatic ranges of each species from the occurrence data, we defined the niche envelop of the two species to contextualize the niche optima of the two species. The range of environmental preferences varied substantially between the two species albeit with visible overlap (CEG: non-parametric environmental gradient Fig. 4A). Interestingly, the peaks of the two niches were far from each other in our environmental gradient analysis although the complete range of the niches showed considerable overlap. The PCAs of climatic niche space also revealed an overlap between the two species, suggesting that marginal populations of both species are adapted to similar environmental conditions (Fig. 4B). Further, when hybrid populations are considered in the analysis, the overlap becomes wider, suggesting niche expansion owing to hybridization (Fig. 4C).

**Figure 4.**
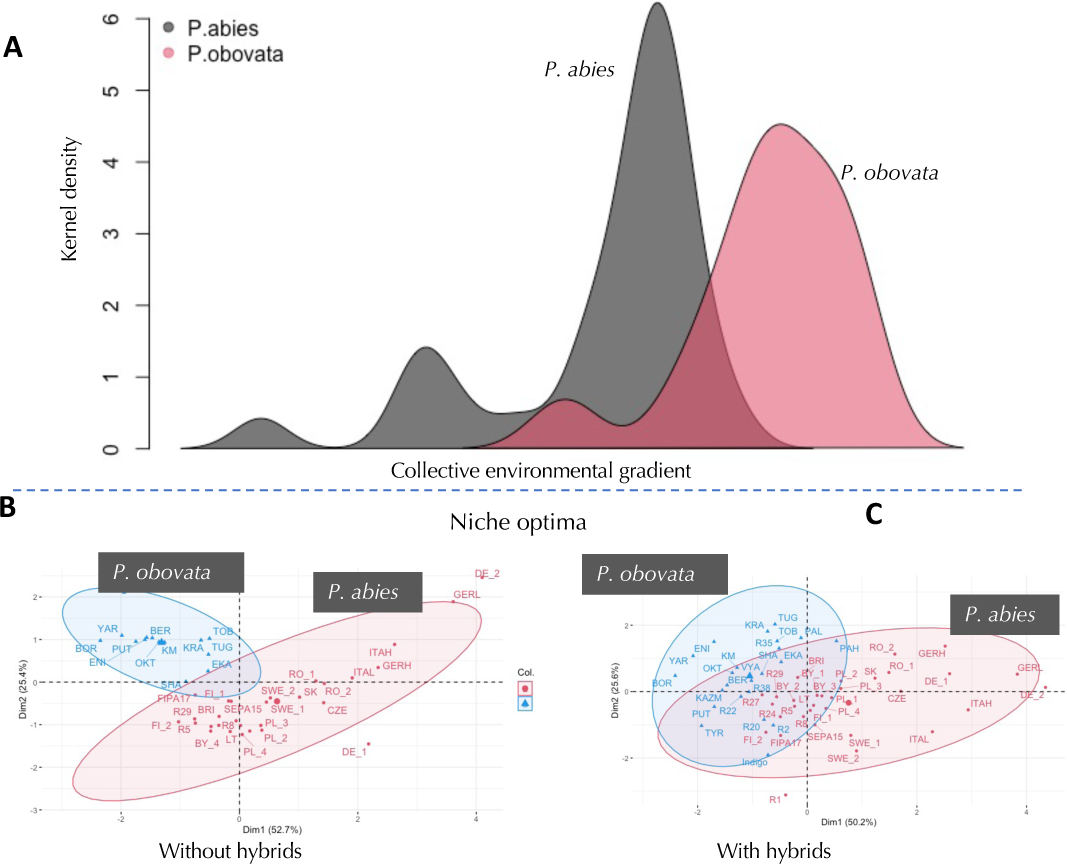
Environmental preference of Norway spruce and Siberian spruce; A. Collective environmental gradient (following Karunarathne et al. 2018): the sum of transformed environmental variables (i.e., Bio8, Bio9, and Bio18) for all populations plotted against Kernel density; B&C. Niche optima of the two species (following Bronnimann et al. 2016).

To shed light on spatial variation in nucleotide diversity, we used niche modeling to infer species’ suitable habitat during LGM (∼25,000 years ago) and mid-Holocene (∼4,000-10,000 years ago Fig. 5). It confirmed that the southern range of *P. abies* (e.g., Alpine, Carpathian, and western European forests) harbored suitable habitat and must have served as refugia during the last glacial maximum. With the warming climate, the species migrated back to higher latitudes, eventually establishing its present distribution. Niche modeling for the period before the Last glacial maximum, specifically for the interglacial period (i.e., ∼140,000 years ago), shows that optimal habitats for *P. abies* existed in the northernmost regions of the current distribution (e.g., northern Sweden and Norway) as well as in the Alpine region. The distribution of Norway spruce thus followed the movement of its niche during cycles of glaciation which drastically affected population connectivity. In contrast, the alteration in suitable habitats for *P. obovata* was considerably more modest across glacial cycles. This species maintained a broader distribution compared to *P. abies*, with an almost uninterrupted range. Notably, the suitable habitat ranges of both species frequently intersected over the course of the glacial cycle, with a notable convergence observed in the western Russian forests during LGM. This overlap could have facilitated multiple instances of hybridization and genetic exchange between the two species. This phenomenon mirrors the current distribution scenario where a modest spatial overlap of the suitable ecological niche is evident in North European Russia (*sensu* TDWG classification) (Fig. 5).

**Figure 5.**
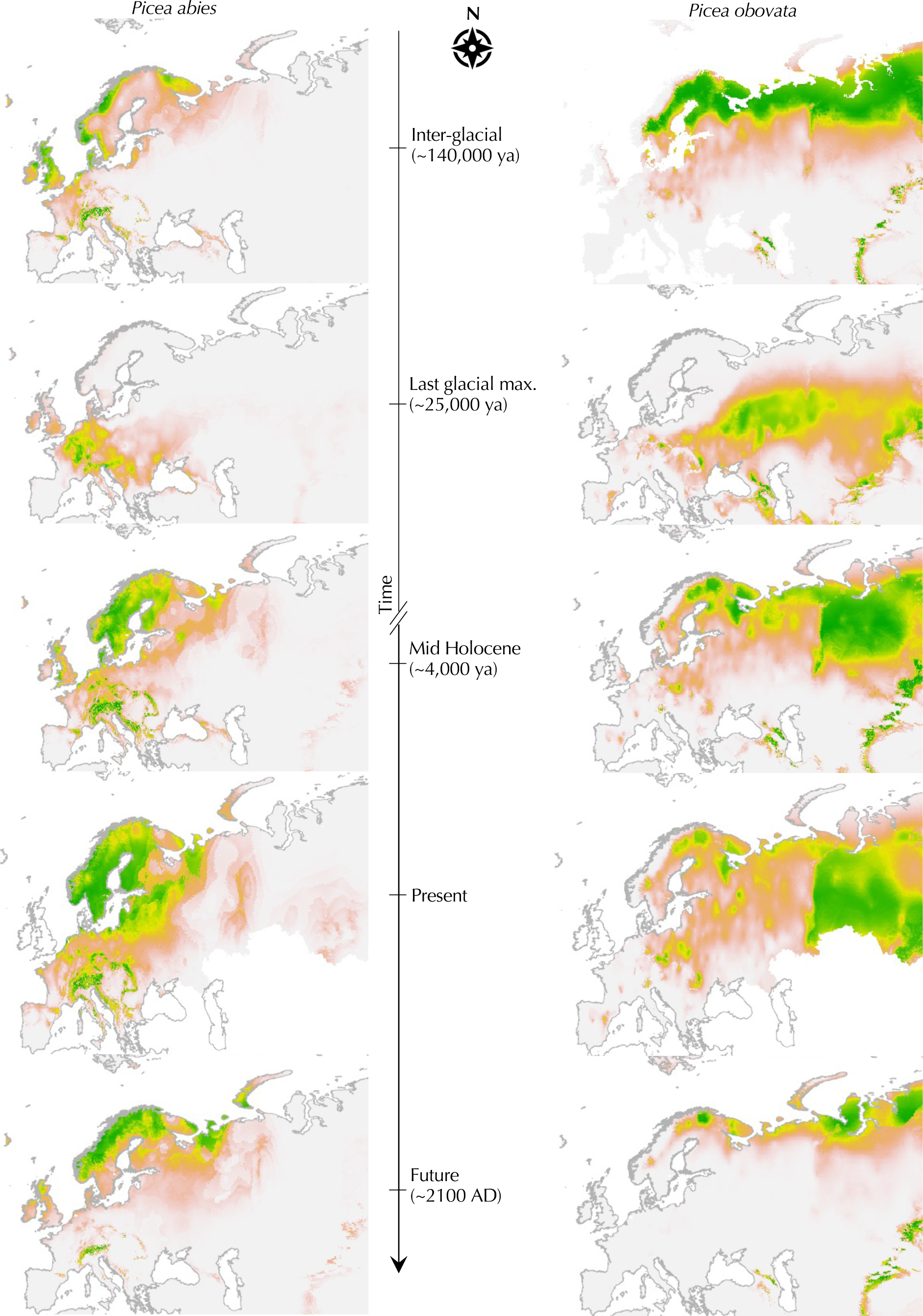
Distribution of suitable habitat/niche for *P. abies* and *P. obovata* from the last interglacial period to future climatic conditions, showing the niche movement from the past to present, determined by maxent modeling of the niche optima.

### Genetic offset and resilience to climate change

Lastly, we explored the capacity of present-day populations of both species to respond to climate change. To achieve this, we quantified population adaptability to future climates by assessing the discrepancy in allele frequencies between current and projected values based on anticipated climatic conditions, referred to as the “genetic offset.” Risk of non-adaptedness (RONA: Restalb et al. 2016) measures the allele frequency change necessary for future climate based on a linear regression of current values. We calculated RONA for three bioclimatic variables using climate projections for the years 2080 and 2100 (i.e., Bio8, Bio9, Bio18). Overall, the RONA assessment indicates a relatively low risk of non-adaptedness to all three bioclimatic variables in regions characterized by high genetic diversity (Fig. 6). This underscores that genetic diversity serves as a pivotal factor in mitigating the risk of non-adaptedness, thereby providing populations with a sheltered wellspring of adaptive potential. We observed high risk of non-adaptedness scores in populations from central Europe and at the Southernmost range of *P. abies* (0.18 – 0.22) indicating that populations in these regions may be at higher risk of extinction under future climatic conditions. However, this trend was not seen in populations in the Alpine region which has served as a refugium during the glacial period and also most likely in warmer conditions during interglacial periods (see Fig. 5).

**Figure 6.**
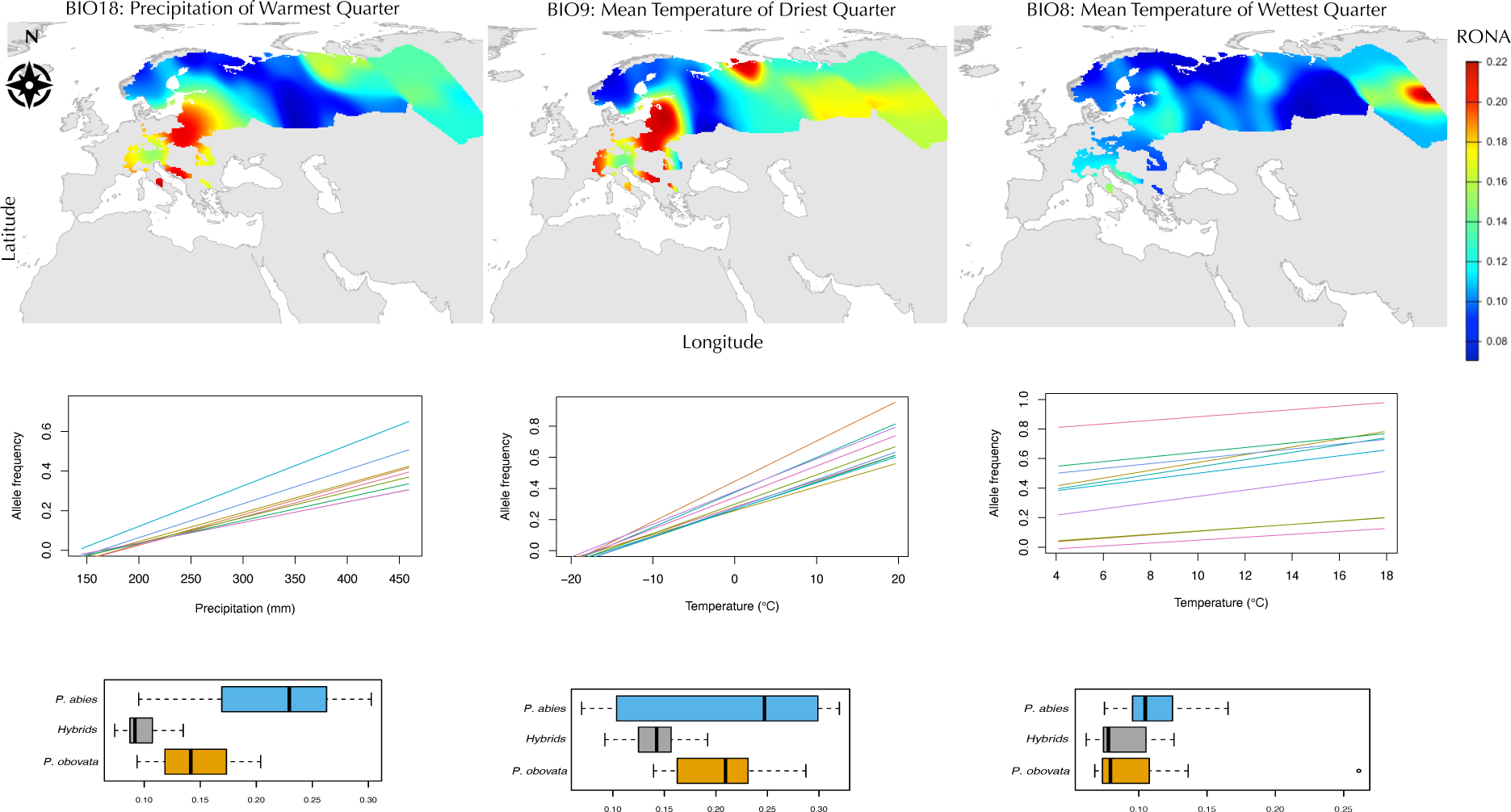
Top row: The map of RONA (Risk of non-adaptedness) for Norway spruce and Siberian spruce. RONA was assessed according to Rellstab et al. 2016, using the coefficient of variation from the linear regression of present environmental variables and allele frequencies. Middle row: Predicted linear response of top 10 alleles to each climate variable. Bottom row: Range of RONA for each bioclimatic variable among the two species and the hybrid populations.

Even more intriguingly, in the case of both species, the risk of non-adaptedness reaches its nadir within the hybrid zones (Fig. 6). This observation further bolsters the proposition that admixture amplifies genetic diversity, thereby fostering the broadening of ecological niche range, a phenomenon conducive to species adaptation to future climatic scenarios. Despite the fact that the Siberian spruce populations harbor the broadest genetic diversity overall (with a π range of 0.0059-0.0069), they paradoxically exhibit a greater susceptibility to non-adaptedness compared to the hybrid populations. Finally, we integrated insights from the genotype-environment-association study with those of the RONA analysis; about two-thirds of the SNPs detected as outliers in the pRDA are among the SNPs with the highest RONA values. This convergence signifies that the SNPs responsible for local adaptation to current climatic conditions are potentially detrimental in the context of long-term response to climate change.

## Discussion

Extensive evidence now highlights the detrimental influence of global climate change on biodiversity at multiple scales (see Altieri & Gedan, 2015; Walther et al., 2002; Williams et al., 2021). Population-level genetic variation equips species with the capacity to navigate shifting climates, safeguarding them from extinction (Meester et al., 2018). This is often observed through species’ adaptations to their local environment and reflects their potential for future climate change resilience (Sang et al., 2022). Nevertheless, identifying these environment-linked alleles necessitates extensive whole-genome sequencing of multiple populations and/or long-running common garden experiments. Nonetheless, this approach proves impractical for species characterized by extended generation times, frequently overlooking essential evolutionary parameters such as demographic history and the extent of gene flow. In the present study, using exome capture sequences, we investigate the role of species hybridization in shaping the current ecological niches of Norway and Siberian spruce. Our findings reveal that recurrent admixture has heightened the climate change resilience and adaptive capacity of both species. Additionally, this approach has enabled us to pinpoint populations vulnerable to prevailing climate change conditions.

### Niche modeling suggests recurrent shifts from Southern to Northern distribution ranges

Our analysis of niche optima movement of Norway spruce during and post-LGM agrees with fossil records and ancient DNA analyses (Brewer et al., 2017; Nota et al., 2022; Parducci et al., 2012; Tollefsrud et al., 2015). Tollefsrud et al. (2015) traced the optimal ecological niches of Norway spruce and Siberian spruce during the most recent glaciation cycle. They also delineated the distribution ranges of these species during the LGM and the mid-Holocene. The study’s findings showcased the gradual migration of both species to and from their southern refugia. This is of course expected since many species in temperate regions followed the same path during the last glaciation cycle (Feliner, 2011; Schönswetter et al., 2005; Sommer & Nadachowski, 2006). However, inferring *P. abies* and *P. obovata* ecological niches further back in time (∼140K years ago), a period where macrofossils fail to provide species-specific information, unveiled a substantial overlap in the distribution ranges of both species in their northern habitats. This convergence could have paved the way for multiple instances of admixture. This is highly consistent with Chen et al. (2019) and Zhou et al. (2023) estimates of a massive admixture event ∼103 - 187 kya between *P. abies* and *P. obovata* populations using genome-wide SNPs. Our niche modeling approach further reveals that the Southern European Mountain ranges, including the Alps and the Carpathians, likely acted as refugia for *P. abies* during both glacial and interglacial periods. Meanwhile, the species’ primary distribution underwent a transition from southern to northern latitudes in tandem with the shifts in glaciation cycles. The recurrent utilization of the same discrete and well-defined refugia across ice-cycles may, at least partially, explain the difference observed between Norway spruce and two other boreal tree species - *Pinus sylvestris* (Scots pine) and *Betula pendula* (Silver birch) – both of which display notably less population structure (Bruxaux et al., 2023; Milesi et al., 2023; Tsuda et al., 2017). The latter may have instead survived the glacial periods in a large metapopulation south of the glaciated regions (Hewitt, 2000; Palmé et al., 2003).

### Genetic diversity does not reflect niche-centrality but captures distribution range variations

Niche centrality suggests that a species’ performance is best at its optimal niche (also niche centroid) conditions (see Lira-Noriega & Manthey, 2014). Although the niche center of a species may or may not be the geographical center of the species, it hypothesizes that the genetic diversity in the niche center is higher than in the peripheral populations (Manel & Holderegger, 2013; Pironon et al., 2015). Our results, however, show little congruence with niche centrality expectations. For *P. abies*, we recognized southern Scandinavia as the niche centroid (Fig. 4: see the populations in the center of eigen ellipses) and populations in this region show the highest nucleotide diversity (Fig. 2A). It is, however, imperative to mention that the currently observed high genetic diversity of *P. abies* in the southern Scandinavian forests is also a result of recent introductions of genetic material from the whole range of the species (Chen et al., 2019 and reference therein). For *P. obovata*, our sampling does not represent the complete current distribution of the species, especially east of the Yenisei River. For the western range of the species, the highest genetic diversity is found along the Yenisei River and the niche centroid points towards the northern Ural Mountains. While acquiring a more continuous sampling would enhance the informativeness of our conclusions, various other factors could also account for the absence of support for niche centrality. First, both Norway and Siberian spruces, as boreal species with intricate demographic histories, experienced shifts in their suitable habitats and ecological niches across multiple ice-age cycles. For *P. abies*, populations were even extirpated during the LGM from what represents today the niche centroid. Additionally, the long generation time and long-range pollen dispersal of coniferous species could also contribute to the lack of fit to the Niche centrality expectations in these species, as is observed in *P. sylvestris* across its whole distribution range (Bruxaux et al. 2023). Our findings align with recent research that observed minimal variation in estimates of genetic diversity among populations from various dominant forest tree species in Western Europe (James et al., 2023; Milesi et al., 2023).

### Recurrent hybridization enlarged the ecological niche breadth

Our niche modeling strategy and retrospective projections offer additional ecological validation for the influence of hybridization on shaping the genetic diversity of both species. Our results are in agreement with genetic data, demographic modeling (Zhou et al. 2023), and paleoecological records (Semerikov et al., 2013 and references therein), which suggest Siberia’s state during the last glacial cycle, and possibly more ancient glacial cycles, as a dry desert interspersed with pockets of forested areas. In agreement with these data, our niche modeling results (Fig. 5) indicate that the suitable habitat of *P. obovata* remained notably larger than that of *P. abies* during the late Pleistocene and Holocene periods. Furthermore, it can be seen in Fig. 5 that the suitable habitat for both species overlapped in many instances during the Pleistocene and Holocene, in particular in their northern range, representing opportunities for genetic introgression. Contrary to limited gene flow, Zhou et al. (2023) demonstrated that secondary contacts among distinct genetic entities of the two species amplified local genetic diversity. This phenomenon is evident in populations situated within present contact zones or within the expansive hybrid zone between *P. abies* and *P. obovata*, which displays high π values compared to neighboring populations (Fig. 2). More importantly, Zhou et al. (2023) highlighted that the interactions between *P. abies* and *P. obovata* led to the emergence of novel genetic clusters with admixed characteristics. These clusters have endured through to the present day, persisting within cryptic refugia during glacial periods. Further, they proposed that during the expansion phases, these entities helped to bridge *P. abies* and *P. obovata* genetic backgrounds, serving as stepping stones (see also Tsuda et al., 2016).

The role of hybridization in allowing niche expansion has been well documented in many cases of invasive species (e.g., Pfennig et al., 2016 and reference therein), particularly in the presence of long-distance gene-flow (e.g., Berthouly-Salazar et al., 2013) that allows for the maintenance of genetic diversity in peripheral populations and for the introgression of favorable alleles from other populations. Hybridization can also directly contribute to favorable traits (e.g., Darwin’s finches’ beak shape, Grant & Grant, 2019) and foster their evolution through an increase in standing genetic variation, while adaptation from *de novo* mutations can be slow (Rieseberg, 1997; Taylor et al., 2015); especially in species with long generation time like spruces.

In Norway and Siberian spruce, primary phenotypic traits related to growth phenology or reproduction exhibit a continuous spectrum of variation between the two species. Hybrid individuals, in this context, manifest intermediate phenotypes (Lagercrantz & Ryman, 1990; Orlova et al., 2020; Popov, 2010). Categorized by their phenotypes, the hybrid populations can be distinguished into two forms, each exhibiting “abies” or “obovata” characteristics (Nakvasina et al., 2019). These distributions align with the delineation of the hybrid zone as outlined by Zhou et al. (2023) and precisely coincide with the distribution of the hybrid index generated in our study (see Fig. 1A). Provenance tests (i.e., breeding experiments equivalent to common garden experiment) also revealed that the genotypes of hybrid nature had higher survival rates in the hybrid zone (Nakvasina et al., 2019). Consistent with these findings, our study demonstrates that the hybrid populations do not necessarily fall within the ecological niches of either species. Instead, they carve out a niche of their own that partially intersects with the ecological preferences of both species (see Fig. 4). Therefore, it is likely that hybridization is responsible for expanding the ecological niche breadth of both species.

### Integrating niche modeling, population genetics, and genetic-offset analysis

Given the current pace of global climate change, the central question is, will long-lived organisms such as spruces have enough time to adapt to the new environmental conditions before facing an inevitable demographic collapse? First and foremost, our niche modeling to future scenarios suggests that suitable habitats would be available for both species in their current northern distribution range and our genetic offset analysis indicates that these populations should be poised to counter climate change (as indicated by a low risk of non-adaptedness, see Fig. 5 and 6). However, it’s worth noting that a considerable portion of the current range of *P. abies*, as well as a substantial segment of the western distribution range of *P. obovata*, might not be suitable for their continued survival.

The populations situated at the opposite ends of the adaptive index, as indicated by the genetic offset analysis, correspondingly represent those populations most susceptible to future climate conditions (Fig. 3A & 6). This observation aligns with the expectation that populations displaying a pronounced degree of adaptedness can be regarded as “specialists” tailored to their specific environments. However, according to our modeling of the future distribution ranges, regions with populations showing an elevated risk of non-adaptedness are projected to accommodate suitable habitats for species persistence (Fig. 5). At first glance, these two results may seem contradictory. The essence of the RONA analysis lies in gauging the average change in allele frequency of SNPs associated with key climatic variables. This change is required to maintain an equivalent level of adaptedness under future climatic scenarios. This suggests that the alleles that are responsible for local adaptation to the current climate might potentially be deleterious under future environmental conditions, and consequently, they could be at risk of being lost. However, RONA, like many other genetic off-set prediction models, suffers from well-identified limitations, including its failure to account for demography history, population structure, or gene flow (Gain et al., 2023; Rellstab et al., 2021).

Both *P. abies* and *P. obovata* exhibit extensive effective population sizes alongside far-reaching pollen dispersal (Zhou et al., 2023). This characteristic could potentially aid in the dissemination of adaptive alleles, particularly considering the distinct ecological preferences of the two species (see Fig. 4). Our study has further revealed that hybrid populations occupy their own ecological niche while having a unique genetic makeup. Moreover, these hybrid populations could serve as potential sources for future recolonizations, in addition to acting as a reservoir of genetic diversity. For instance, hybrid populations display the highest survival rate within the hybrid zone (Nakvasina et al., 2019). Both species hold significant economic importance, and human-assisted transfer of trees from different provenances could play a role in bolstering adaptation to future conditions. For instance, *P. abies* trees of Southern origin planted in Southern Scandinavia for breeding purposes (J. Chen et al., 2019) exhibited higher growth rates in Southern Sweden compared to trees of local origin (Milesi et al., 2019). Intriguingly, Southern Sweden presently showcases the highest genetic diversity in *P. abies* (Fig. 2A). Consequently, these trees can potentially introgress with local counterparts leading to the introduction of favorable alleles for future climatic conditions. Notably, the plants’ response to global change can be rapid, even in long-lived species, like oak (*Quercus petraea*), which despite its ∼50-year generation time, displayed a genome-wide response to the Little Ice Age (∼1650 AD, Saleh et al., 2022).

Finally, while neither of the studied spruce species might face a large risk of global extinction under future climatic conditions, it is worth noting that certain areas are under particular threat. Among populations with the highest risk of non-adaptedness, two are located south of the Italian Alps and one, Indigo, in the Northern range of *P. obovata* western domain. As highlighted by Zhou et al. (2023), these populations are geographically isolated from the rest, exhibit low nucleotide diversity, and have undergone more pronounced genetic drift. These characteristics often point to populations that might struggle to contend with the challenges posed by global climate change.

## Conclusion and Perspective

When species are unable to track their ecological niche, standing genetic variation is the main determinant for species’ ability to cope with climate change. This is even more true for species with long-generation time where the probability of adaptation through *de novo* mutation is low. Cycles of range contractions and expansions can enhance genetic diversity in a feedback loop (Barrett & Schluter, 2008; Sexton et al., 2009) albeit with expected diversity loss during contraction events (Pauls et al., 2013). In Western Europe, species-wise genetic diversity of seven forest tree species globally increased across multiple ice-cycles (Milesi et al., 2023). Among them, at least five (*Betula pendula*, *Quercus petraea*, *Fagus sylvatica*, *Populus nigra, and P. abies*) are known to hybridize with closely related species and are all long-lived species with long-range gene flow. Hybridization could thus be a main factor in coping with climate change in temperate and boreal forest tree species, as our study suggests in the *P.abies* / *P. obovata* species complex. There is a burst of new methodology development aiming at integrating niche modeling and genetic offset prediction in order to better apprehend species response to global change (e.g., Barratt et al., 2023; Y. Chen et al., 2022). If this clearly represents a step forward, our study also evidenced the need for a more integrative framework to also consider species demography history, population structure, and gene flow.

## Materials and methods

### Sampling, genetic material, and SNP calling

The populations come from a large-scale study uncovering the demography history and spatial population structure of the closely related Norway spruce (*Picea abies*) and Siberian spruce (*Picea obovata*) species (Zhou et al., 2023). DNA was extracted from the needles of 546 individual trees coming from 55 populations representing five predetermined geographic groups (including introgress individuals and putative hybrid populations: following (J. Chen et al., 2019; L. Li et al., 2022; Nakvasina et al., 2019); see map Fig. 1) and genomic DNA was extracted for NextGen sequencing using Qiagen® plant DNAeasy kit. Paired-end Illumina libraries were constructed and sequenced by RAPiD Genomic, USA, using 40,018 exome capture probes targeting 26,219 genes (Vidalis et al., 2018). Raw reads can be found on SRA (https://www.ncbi.nlm.nih.gov/sra) under PRJNA511374 and PRJNA1007582.

Clean sequence reads were aligned to the *P. abies* reference genome (Nystedt et al., 2013) with BWA (H. Li & Durbin, 2009), and individual genotypes were identified using GATK HaplotypeCaller (Van der Auwera & O’Connor, 2020) followed by SNP calling across all samples with GenotypeGVCFs and hard-filtering of SNPs (QD < 2.0, MQ < 40.0, SOR > 3.0, QUAL < 20.0, MQRankSum < -12.5, ReadPosRankSum < -8.0, and FS > 60.0). For population genetics analyses, we retained only putatively neutral SNPs (i.e. synonymous mutations, mutations located in the introns or into the intergenic space) and pruned the dataset for singleton and SNPs in high linkage disequilibrium (plink v1.9, www.cog-genomics.org/plink/1.9/, pairwise *r^2^* > 0.2, window size 1 kb), resulting in a 390,903 SNPs data set. For all other analyses, we kept all SNPs with minor allele frequency higher than 0.05 including both neutral and non-neutral alleles (450,000 SNPs).

### Climatic data selection

Environmental variables for current conditions were downloaded from the WorldClim 2.0 database at 30 arc seconds resolution (Fick & Hijmans, 2017; Hijmans et al., 2005). They are annual averages of both temperature and precipitation-related variables. Climatic records at population locations were extracted with the TERRA R package (Hijmans et al., 2022). In order to avoid multi-collinearity and model overfitting, we performed pairwise Pearson-correlation between all the bioclimatic variables, sequentially removing the variable with the highest correlation coefficient, *r*, until it reached *r_max_* = 0.6. Additionally, to ensure that multi-collinearity is minimal in the composite variables of RDA, we assessed the variance inflation factor (VIF) and retained the environmental variables for which the VIF was lower than five, as recommended by Capblancq & Forester (2021). We used this final set of environmental variables (predictor variables hereafter) for all downstream analyses.

### Hybrid index

We calculated a *hybrid index* using the ADMIXTURE software (v.1.3.0 Alexander et al. 2009) assuming *K* = 2 ancestral populations. The hybrid index was defined as being the proportion of *P. abies* ancestry component and ranges from 0 “pure” *P. obovata* to 1 “pure” *P. abies*. The population-wise *hybrid index* was obtained by averaging individual *hybrid index.* Populations with a *hybrid index* higher than 0.2 and below 0.8 were classified as hybrid populations in qualitative analysis. Extrapolation of the *hybrid index* to the whole distribution of the two species was done using custom-R scripts (available in the [GitHub](https://github.com/piyalkarum/RC_EAA) repository) with nearest neighbor interpolation and was mapped to visualize the spatial distribution of genetic diversity of the two species.

We calculated global nucleotide diversity (*π*: Nei and Li, 1979) to assess population-level genetic diversity with the POPGENOME R package (Pfeifer et al., 2014). Extrapolation of the *π* values to the whole distribution of the two species was done using the same approach as for the *hybrid index.* We then investigated the relationship between the *hybrid index* and various metrics of genetic diversity and variation in allele frequencies. In order to minimize the skewed allele frequencies due to the varying number of samples in each population, we (i) randomly sampled ten individuals per population for populations with more than ten individuals and (ii) grouped close-by populations that were also genetically similar before downsampling to ten individuals. The resulting dataset consisted of thirty-six populations.

### Genetically informed climatic group definition

In order to find genetically informed “climatic groups” of both species, we combined two sets of data: i) adaptively enriched genetic space scores from pRDA – the RDA scores represent local adaptation in populations derived from allele frequencies, ii) bioclimatic data – this dataset represents the environmental variability of populations after removing of multi-collinearity. The former was used as a proxy for population-level adaptations to local climatic conditions and the latter to explain the spatial variation of local conditions. A *k-means* clustering was performed for the total within the sum of squares to determine the optimal number of genetically informed climatic clines. This was performed separately for tentative hybrid and non-hybrid populations to avoid cluster overlaps that could arise from specific adaptations in hybrid populations. In non-hybrid populations, we identified five distinct climatic groups while there were only three within putatively hybrid populations. After obtaining the optimal number of clusters as five, a second k-means clustering was performed with K=5 for all populations (i.e., including hybrid and non-hybrid populations) to determine the climatic group membership of each population. This step was performed specifically to group the putative hybrid populations with their closest non-hybrid populations to identify their closest climatic groups.

### Identification of alleles putatively involved in local adaptation

To study the genetic architecture of local adaptation in Norway spruce and Siberian spruce, we ran genome scans to detect outlier-loci (i.e., loci showing an excess of genetic differentiation in comparison to the rest of the genetic background) using the principal component-based outlier detection method, “pcadapt”, implemented in the R package “pcadapt” v.4.3.2 (Luu et al., 2017; Privé et al., 2020). Briefly, a Z-score matrix, obtained from regressing all SNPs to PCs, is used to produce one multivariate distance for each variant. Assuming that SNP variation along principal component axes largely reflects demographic processes, this method allows controlling for population structure when testing for local adaptation. In our analysis, to control for false positive risk discovery associated with multiple testing, we used FDR q-value < 0.1 to obtain the significant SNPs putatively involved in local adaptation.

Redundancy Analysis (RDA) is an asymmetric canonical analysis that combines ordination and regression, where multiple regression on the response variable (i.e., allele frequencies) and explanatory variables (climatic variables) is followed by ordination of fitted values of the regression (Van Den Wollenberg 1977). We used Redundancy Analysis (RDA) to find the main climatic variables that explain most of the variance in allele frequencies between the two species and to identify candidate loci associated with environmental variation.

In the special case of partial RDA (pRDA), additional explanatory variables (covariates) are used to control for various confounding effects (see Capblancq et al., 2018; Capblancq & Forester, 2021). This aspect of RDA is extremely useful in association analyses to control for the confounding effect of factors that are not the focus of the study, such as population structure and geographic distance. Thus, to isolate different drivers of genetic variation, in addition to a full model (including all predictor variables), we performed three independent pRDA models controlling for geographic distance, population structure, and environmental conditions. We adopted the method described in Capblancq & Forester (2021) where the Mahalanobis distance of each locus to the center of the RDA is calculated using two axes (K=2) of partial RDA (i.e., the number of axes that explain more than 60% variation; following Capblancq et al. 2021), followed by obtaining the top outlier SNPs from this distance matrix. For geographic distance, the cartesian distance of populations was used as a proxy; to control for population structure, we used the first three axes of a PCA performed on the set of putatively neutral loci.

Using the output from the pRDA, we computed an *Adaptive index (Aij)* as implemented by Capblancq & Forester (2021) from Steane et al. (2014) using the RDA 1 and 2 independently. For the top outliers, RDA1 and RDA2 explained 88% and 10% of the variation respectively. For each location, *j*, the adaptive index is given by the sum over each climatic variable, *i*, of the products between its score along a given RDA axis and its value at location *j*.

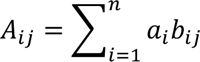

*A_ij_* ranges from negative to positive values and essentially measures the match between the genotypes and their environment, 0 meaning no match.

### Niche expansion, species distribution modeling, and historical niche optima movement analysis

The climatic niche of the two species was defined as the ordination space of all the populations with PCA eigenvalues. Therein, a PCA was performed on the selected climatic variables (i.e., Bio8, Bio9, Bio18) for all the populations of *P. abies* and *P. obovata* separately. The optimal niche was determined as the centroid of the eigenvalues and the niche breath was defined as the 95% ellipsoid of the PCA scorings. The analysis was performed with and without hybrid populations to visualize the niche breadth overlap. As a complementary analysis, we also used a non-parametric method, collective environmental gradient (CEG; Karunarathne et al. 2018) to assess the total environmental aptitude of each species and to visualize their overlap. CEG is a single-value decomposition approach to determine the composite two-dimensional environmental/niche breach. We performed CEG according to Karunarathne et al., (2018) for the selected bioclimatic variables (i.e., Bio8, Bio9, Bio18).

Species distribution models for the current, mid-Holocene (ca. 6,000 ya), Last Glacial Maximum (LGM, ca. 22,000 ya), Last Inter-glacial period (ca. 120,000 - 140,000 ya), and future climatic scenarios (2021-2080 and 2081-2100) were constructed using maximum entropy (Maxent) modeling (Phillips et al., 2006). The optimal niche for separate species was determined with selected climatic variables (i.e., Bio8, Bio9, Bio18) using the *maxent* function of the *dismo* R package (Hijmans et al., 2016). Maxent uses presence-only data to estimate the probability distribution of occurrence based on environmental constraints (Phillips et al., 2006). In order to test the models and to facilitate a more accurate prediction of species distribution, we downloaded species occurrence data for *P. abies* and *P. obovata* from GBIF (www.GBIF.org) and filtered the data to include only confirmed and natural occurrences of the species. occurrence data were then subsampled to retain only one occurrence per grid cell of 1 arc minute. The final dataset consisted of 619 occurrences (*P. abies*: 288; *P. obovata*: 331), hybrid populations have been excluded. All bioclimatic variables were downloaded from WorlClim 1.4. database at 30 arc-second resolution. The climatic variables for the Mid-Holocene, LGM, and Last inter-glacial were downloaded from the model CCSM4 and the future climate variables were downloaded for ACCESS-CM2 model. The projection of past and future niche distribution of the two species was done using the *predict* function of the *dismo* R package. All the models were evaluated using the area under the operating curve (AUC) and only the models with an AUC of at least 0.85 were considered.

### Assessment of genetic offset and resilience to future climatic conditions

The genetic offset of an entity (e.g., population) is the allele frequency difference between the current and required frequency in the future to survive in future climatic scenarios. We calculated the genetic offset of our populations with risk of nonadaptedness (RONA) as in Rellstab et al. (2016). RONA is the average theoretical change in allele frequency at climate-associated loci required to match future climatic scenarios. This is established by regressing the selected loci allele frequencies with the selected environmental variables and using the regression coefficients to predict the expected allele frequencies of the respective loci to future conditions. The difference between the two allele frequencies is the risk of nonadaptedness. Here, we first performed linear regression on all loci with the selected environmental variables (i.e., Bio8, Bio9, Bio18) and kept only the loci that showed significant association (*p*-value < 0.05) with the three environmental variables. Then we obtained the intersection between these alleles and the top outliers from the RDA analysis for the RONA analysis. To avoid linkage between selected loci, we filtered loci within 10,000 base pairs in the same scaffold; the locus with the highest R^2^ from the regression was retained. In total, ∼25,000 candidate SNPs for environmental associations were kept for the analysis. Finally, we used nearest neighbor interpolation (as in Rukundo & Cao 2012: custom scripts available in the [GitHub](https://github.com/piyalkarum/RC_EAA) repository) to predict the RONA values for the entire distribution of the two species for spatial visualization (Fig. 6).

## Acknowledgments

We would like to thank Elena Nakvasina, Vladimir Semerikov, Lars Opgenoorth, Katrin Heer, and Giovanni Giuseppe Vendramin for their valuable efforts in providing us with plant materials. We also thank the Swedish National Infrastructure for Computing (SNIC) for resource allocation for high-power computing under the project numbers SNIC 2021/6-313 and SNIC 2021/5-540. The Nilsson-Ehle grant by the Royal Physiographic Society of Lund awarded to Piyal Karunarathne (42191) was immensely helpful for acquiring genome sequences to fill data gaps.

## Data availability

All the raw reads studied in this study can be found on the NCBI SRA database (https://www.ncbi.nlm.nih.gov/sra) under PRJNA511374 and PRJNA1007582. The scripts used for all the analyses, simulations and respective input data are available in the [GitHub](https://github.com/piyalkarum/RC_EAA) repository.

## Conflict of interest statement

The authors declare no conflict of interest.

## Notes

https://github.com/piyalkarum/RC_EAA

### Competing Interest Statement

The authors have declared no competing interest.

### Summary of Updates

A new hybridization index analysis was done. The niche expansion analysis was redone. The structure of the manuscript was changed.

