## Supplementary material for "Hybridization Mediated Range Expansion and Climate Change Resilience in Two Keystone Tree Species of Boreal Forests": supp.info.


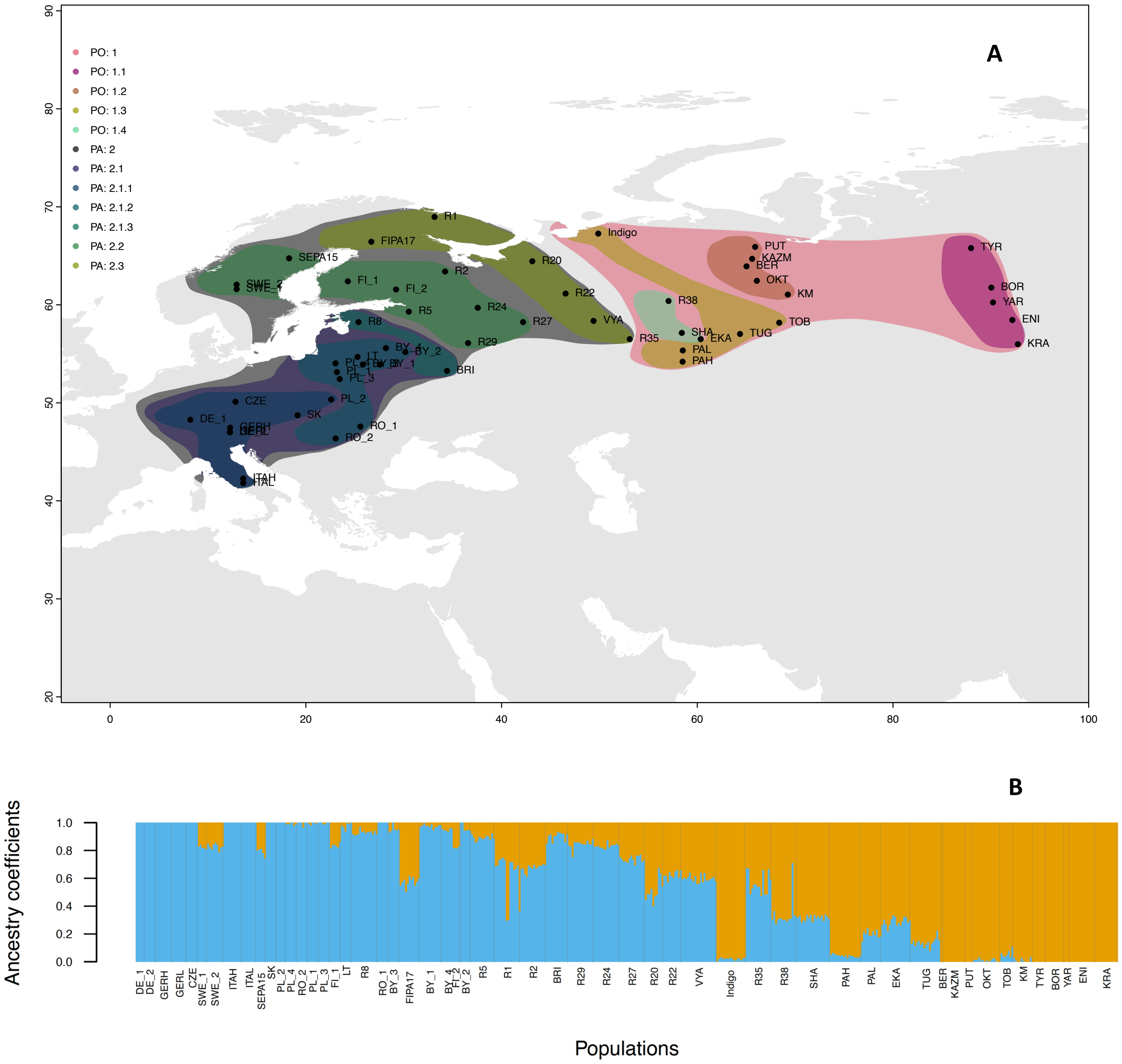


Fig. S1. A. The map depicting the distribution of clusters identified in the genetic clustering (i.e., Bayesian analysis) by Zhou et al. 2023 – PO1 separates all *P. obovata* populations, PA2 includes all the *P. abies* populations; the subclusters within each major cluster is indicated in different colors, the labels on the map shows the population names as in figure 1A**. B.** Ancestry coefficient estimated using ADMIXTURE software and K = 2, blue represents *P. abies* genetic background and orange *P. obovata*.

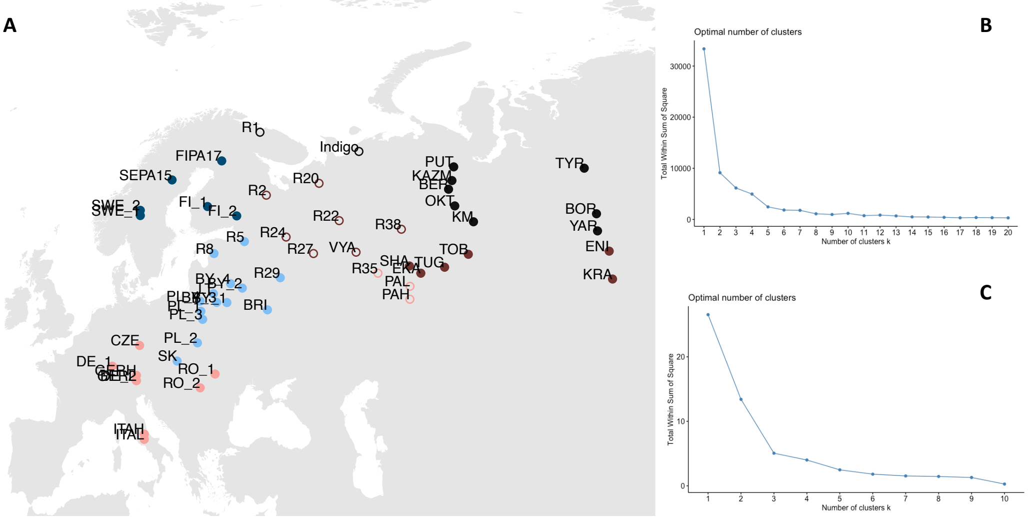


Fig. S2. Clustering of Norway spruce and Siberian spruce based on genetically conditioned environmental variation (climatic clines); A. Distribution of different climatic clines, B. Optimal clusters of all populations, C. Optimal clustering for only hybrid populations

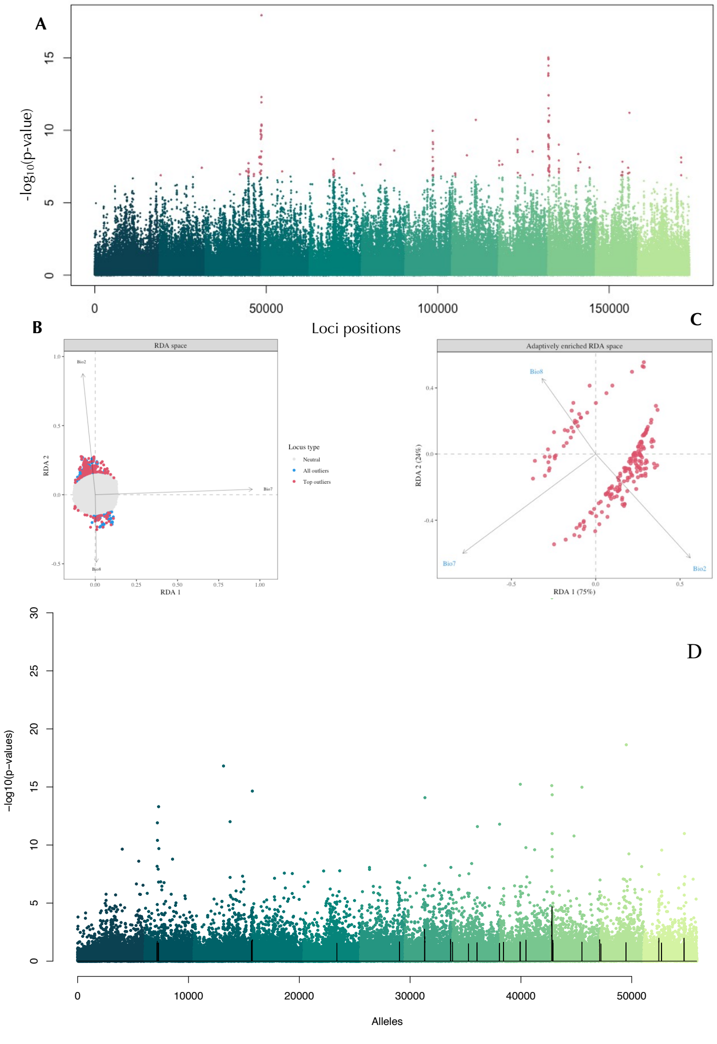

Fig. S3. The enrichment of outlier alleles associated to temperature variables; A: The significance values (-log10(p-values)) from the linear regression of allele frequencies and the environmental variables in RDA laid along the genome. The different shades of green color show linkage groups and the red points are the top outliers; B&C: Ordination of adaptive alleles with the respective environmental variables; D. Output of PCAdapt showing the structuring of loci under selection along the genome; the black line shows major peaks, the different shades of green shows pseudo-chromosomes.

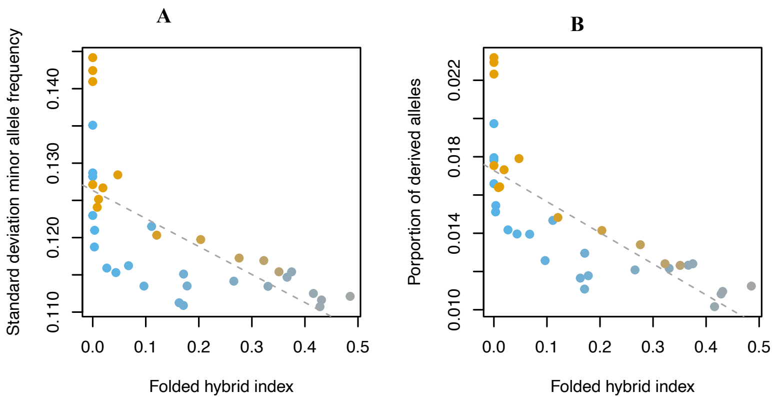


Fig. S4. Relationships between various genetic diversity metrics and hybrid index. The hybrid index has been folded and varies from 0 (“pure” *P. abies* or *P. obovata*) to 0.5 “pure” hybrid populations. Colors gradients reflect the genetic background from “pure” *P. abies* in blue to “pure” *P. obovata* in orange. The gray dotted lines represent the linear regression between the corresponding factors, all $F$-values are significant $p < 0.001$.
